# Transcriptomic networks implicate neuronal energetic abnormalities in three mouse models harboring autism and schizophrenia-associated mutations

**DOI:** 10.1101/625368

**Authors:** Aaron Gordon, Annika Grønborg-Forsingdal, Ib Vestergaard Klewe, Jacob Nielsen, Michael Didriksen, Thomas Werge, Daniel Geschwind

## Abstract

Genetic risk for psychiatric illness is complex, so identification of shared molecular pathways where distinct forms of genetic risk might coincide is of substantial interest. A growing body of genetic and genomic studies suggest that such shared molecular pathways exist across disorders with different clinical presentations, such as schizophrenia and autism spectrum disorder (ASD). But how this relates to specific genetic risk factors is unknown. Further, whether some of the molecular changes identified in brain relate to potentially confounding antemortem or post-mortem factors is difficult to prove. We analyzed the transcriptome from the cortex and hippocampus of three mouse lines modeling human copy number variants (CNVs) associated with schizophrenia and ASD: Df(h15q13)/+, Df(h22q11)/+, and Df(h1q21)/+ which carry the 15q13.3 deletion, 22q11.2 deletion, and 1q21.1 deletion, respectively. Although we found very little overlap of differential expression at the level of individual genes, gene network analysis identified two modules of co-expressed genes that were dysregulated across all three mouse models. One module observed in both cortex and hippocampus was associated with neuronal energetics and firing rate, and overlapped with changes identified in post mortem human brain from SCZ and ASD patients. These data highlight aspects of convergent gene expression in mouse models harboring major risk alleles, and strengthen the connection between neuronal energetic dysfunction and neuropsychiatric disorders in humans.

## Introduction

The complex and highly polygenic nature of psychiatric disorders ^1, 2^ makes it difficult to connect genetic risk factors to brain function ^3^. One way of addressing this issue is by collecting large sample sizes that can capture patterns of convergence among many individuals with the disorders. For example, transcriptomic studies in humans suggest that despite having diverse genetic etiologies and no gross pathology, patients with major psychiatric disorders have recognizable, partially overlapping patterns of dysregulation in gene expression ^4–7^.

Another avenue of research is to focus on the rare, but relatively highly penetrate forms of these disorders ^3^. In severe mental disorders, including in schizophrenia (SCZ), an important subset of these rare highly penetrate forms^8^ is copy number variation (CNV)^2^. Published studies indicate that between 2.5 to 5% of SCZ patients carry one of the eight CNVs most frequently associated with SCZ ^2^. These CNVs have relatively high penetrance compared with common variants and have clearly defined and evolutionary conserved genomic structure^2, 9^, which makes them well suited for generating mouse models to study the brain mechanisms underlying neuropsychiatric disease risk ^9–11^. Additionally, all of these CNVs have pleotropic effects, variably increasing risk for a range of neurodevelopmental outcomes, including intellectual disability (ID), autism spectrum disorder (ASD) and SCZ. Such pleiotropy has been the rule, rather than the exception for most major effect risk genes ^3, 12^. Transcriptomic studies of patients with major psychiatric disorders have largely been based on cases without defined genetic etiologies. Therefore, with the exception of (dup)15q11-13 syndrome and ASD ^6^, it is not known whether specific genetic forms of mental disorders show similar patterns of convergence.

To address whether specific genetic variants imparting major risk for schizophrenia or ASD converge at the molecular level, we leveraged three previously published mouse models with CNVs carrying significant risk for these disorders, Df(h15q13)/+ ^13–15^, Df(h22q11)/+ ^16^, and Df(h1q21)/+ ^17^, which harbor the 15q13.3 deletion^18^, 22q11.2 deletion ^19^, and 1q21.1 deletion^2^ respectively. These three models have strong construct validity as they all harbor deletions of regions homologous to those found in human deletions associated with SCZ and ASD ^2, 9, 19^. Of note, these mice were tested for face validity with respect to phenotypic aspects of SCZ and ASD. For example, Df(h1q21)/+ mice display abnormalities in the dopaminergic pathway^17^ which has been implicated in SCZ ^20, 21^. Both Df(h15q13)-/- and Df(h15q13)/+ mice, which have qualitatively similar phenotypes^9, 13^ displayed phenotypes analogous to some of the negative symptoms and electrophysiological abnormalities similar to changes seen in SCZ patients^13–15^. Additionally, these mice displayed altered social behavior similar to changes seen in ASD patients^13–15^. Df(h22q11)/+ mice displayed a decrease in pre-pulse inhibition and displayed acquired acoustic startle response similar to that seen in SCZ patients ^16, 22^, which could not be rescued by the anti-psychotic drugs haloperidol or clozapine^16^ commonly used to treat SCZ ^23^. While the three mouse models all present neuropsychiatric disease relevant phenotypes, there is substantial phenotypic variability between them, consistent with the individuals carrying these CNVs who present with different diagnoses ^24–26^.

In this study we aimed to explore whether despite the variability in phenotype there was convergence at the molecular level. The variable phenotype in SCZ and other psychiatric disorders does not preclude shared molecular pathways, as many psychiatric disorders show substantial overlap in molecular pathways ^4, 5^. To this end, we performed genome-wide transcriptome sequencing on both the cortex and hippocampus in adult animals from these three mouse models. We discovered many significantly differentially expressed genes in all tissues examined, but found very little overlap between models at the level of standard single gene analyses, even between hippocampus and cortex from the same CNV model. However, examining coordinated transcriptional regulation via gene network analysis revealed modules of genes that were dysregulated across all mouse models. This includes a cortical module associated with mitochondrial function in neurons that was down-regulated across all three mouse models. This analysis is the first to show a convergent pattern of gene expression in mouse models harboring mutations increasing risk for SCZ and allied neurodevelopmental disorders, further strengthening the hypothesis that neuronal energetic dysfunction plays a role in SCZ ^27^.

## Methods

### Samples

Mice were bred and housed until 8 weeks by Taconic MB (Lille Skensved, Denmark), and then transported to the Lundbeck animal facility. Detailed description of animal housing and handling has been previously described ^15^. Adult mice (>10 weeks old) harboring the CNV as well control littermates were sacrificed by cervical dislocation and parietal cortex or hippocampus were dissected by hand and stored at −80°C. With the exception of Df(h15q13)-/- the cortex and hippocampus were dissected from separate animals. All studies were carried out in accordance with Danish legislation, granted by the animal welfare committee, appointed by the Danish Ministry of Food, Agriculture and Fisheries-Danish Veterinary and Food Administration. Samples were lysed and homogenized in RA1 lysis buffer with 1% β-mercaptoethanol (Macherey-Nagel, Düren, Germany) using a PT 1200 E Polytron® (Kinematica, Luzern, Switzerland). Total RNA was purified using TruSeq® Stranded mRNA Sample Preparation kit according to manufacturer’s instructions (Illumina, San Diego, Ca, USA). RNA quality was assessed by measuring RNA integrity (RIN) values and absorption spectra on an Agilent bioanalyzer (Agilent Technologies, Santa Clara, CA, USA). 3’ poly-adenylation purification, adapter ligation, and PCR amplification was carried out using the TruSeq® Stranded mRNA Sample Preparation kit per manufacturer’s instructions, with a minimum amount of 160 ng RNA per sample (Illumina).

### Sequencing

Samples were multiplexed and genotypes were balanced on lanes and flow cells per study. Samples were sequenced in 50 bp single reads or 100 bp paired reads on Illumina HiSeq (Illumina). See sequencing details for each dataset in Supplementary Table 1. Sequencing quality was assessed using fastQC (http://www.bioinformatics.babraham.ac.uk/projects/fastqc/). Reads were mapped to GRCm38 using Gencode v11 via STAR ^28^. Gene expression levels were quantified using the featureCounts function from subread ^29^. Alignment, GC bias and duplication metrics were collected using Picard tools (http://broadinstitute.github.io/picard/) functions CollectRnaSeqMetrics, CollectGcBiasMetrics, and MarkDuplicates respectively and were then aggregated using MultiQC ^30^. Genes with less than 10 reads in over 80% of the samples were dropped. Samples were then split by study and outlying samples were defined as having standardized sample network connectivity Z scores <-2 ^31^, and were removed. All code used for the analyses is available at https://github.com/dhglab/mouse-models-harboring-CNVs.

### Differential Gene Expression

Differential gene expression was performed on each tissue from each mouse line separately. Expression values were normalized using the trimmed mean of M-values (TMM) method from the edgeR package ^32^ followed by voom from the limma package ^33^. Differentially expressed genes were then defined as having a p value < 0.005 using the limma package ^33^, a threshold that we and others have previously validated ^34, 35^. The model used for the differential expression included the following covariates: ∼ Genotype + SeqPC1-SeqPC5. SeqPCs are the first 5 principal components calculated from the Picard sequencing statistics and are used to control for technical variation introduced during sequencing. For one of the dataset (labeled study 2) only 4 PCs were used due to available degrees of freedom.

### Weighted Gene Co-Expression Network Analysis

Hemizygous samples and their matching controls were used to construct a gene co-expression network. The sequencing covariates were first regressed out of the expression data. The CNV was also regressed out while retaining the information regarding the genotype (wt/hemizygous) of the animal to allow us to study common perturbations across all CNVs. The network was constructed using the WGCNA package ^36^ in R for each tissue individually. A soft threshold was chosen to achieve approximate scale-free topology (R^2^>0.8) ^36^ resulting in a soft-threshold of 4 for the cortical network and 5 for the hippocampal network. The network was then constructed using a topological overlap dissimilarity matrix. A dendrogram was created using hierarchical clustering and the modules were defined as branches of this dendrogram using the hybrid dynamic tree-cutting method ^37^. To relate these modules to traits, the first principal component of the module (the module eigengene) was related to genotype (wt/hemizygous), using a linear model. Modules were considered to be associated with the phenotype when fdr < 0.1. Module hub genes were defined as being those most highly correlated to the module eigen gene (kME > 0.7). The top 15 hub genes are plotted along with their top intra-connections.

### Rank-rank hypergeometric test

We performed a rank-rank hypergeometric test as previously described ^38^. Genes were ranked according to their logFC before running the rank-rank hypergeometric overlap test to evaluate overlap between two datasets.

### Go-term enrichment

Gene ontology analysis was performed used GO-Elite ^39^ with default settings and 20,000 permutations. The top 5 biological process and molecular function categories were plotted selected from categories containing at least 20 genes of which at least 5 gene overlapping the test data set. The denominator for the enrichment was all expressed genes in the tissue (all genes passing the filtering cutoff from the tissue).

### Cell type enrichment

Cell type enrichment was performed using a Fisher exact test using data sorted cells ^40^. P values were corrected for multiple testing using FDR. For single cell enrichment the Expression Weighted Cell Type Enrichment (EWCE, https://github.com/NathanSkene/EWCE)^41^ was used in conjunction with single cell data from mouse cortex and hippocampus ^42^. EWCE was run with 10,000 bootstrap repetitions on both levels of cell types.

### GWAS enrichment

GWAS enrichment was performed as previously described ^5^ on the cortical modules significantly associated with genotype (wt/hemizygous). To test for enrichment, a stratified LD score regression ^43^ was used. SNPs were assigned to a module if they were within 10 kb of a gene in the module. Enrichment was calculated as the proportion of SNP heritability accounted for by each module divided by the proportion of total SNPs within the module. Significance was tested using a jackknife method, followed by FDR correction. Enrichment was tested in GWAS from ASD^44^, SCZ ^45^, BD ^1^, and MDD ^46^.

### Firing rate

Neurons were grouped into two groups, slow and fast spiking neurons, based on Sugino et al 2006 ^47^. The cM2 module eigengene was calculated for each of the cell types based on microarray data from the same study (GSE2882)^47^. The two groups of neurons were compared using a two-sided Mann–Whitney–Wilcoxon test ^48^.

### Protein-protein interactions

Protein-protein interactions were mapped using the Dapple ^49^ module version 0.17 in GenePattern (http://genepattern.broadinstitute.org) with default settings.

### Module preservation

Module preservation was performed to test whether the modules identified in the hemizygous mice were also found in the homozygous 15q13.3 deletion mice. A Z-summary statistic that aggregates many preservation measurements ^50^, based on 200 permutations, was used to determine preservation. Modules with 2 < Z_summary_ < 10 were considered moderately preserved while modules with Z_summary_ > 10 were considered strongly preserved. Modules with Z_summary_ < 2 were not considered to be preserved.

### Mean Gene rank test

The mean gene rank test was performed as previously described ^34^. For each gene set we measured the average ranking of the differentially expressed genes in all the data sets ranked by their p-values. The average ranking was normalized as follows:

*Score=1-Mean ranking of Differentially expressed genes /Total number of genes* such that if all DEG were at the top of the list the score would approach 1 and a random distribution of them throughout the list would results in score of 0.5. A p-value was calculated using 10,000 permutations which was then corrected for multiple testing using the Benjamini-Hochberg false discovery rate ^51^.

### Data availability

Code and data used for DE and for the construction of gene networks can be found at https://github.com/dhglab/mouse-models-harboring-CNVs. Data is available at the NCBI Gene Expression Omnibus (GEO), accession number GSE129891.

## Results

### Differential gene expression in the CNV/tissue combinations

We sequenced the transcriptome of both the cortex and hippocampus from three different mouse lines with hemizygous deletions modeling human CNVs Df(h22q11)/+ ^16^, Df(h1q21)/+ ^17^, and Df(h15q13)/+ ^13^ as well as mice carrying the homozygous deletion Df(h15q13)−/− ^15^. These mice model the 22q11.2, 1q21.1 and 15q13.3 human CNVs respectively ^13, 15{, 16, 17^. We analyzed differential gene expression in each CNV/tissue combination separately (Figure 1A), comparing the transgenic mice to matched littermate controls. The magnitude of differential expression (DE) genes was similar between the hemizygous mice harboring the 3 different CNVs (84 and 135 in Df(h22q11)/+ cortex and hippocampus, 199 and 26 in Df(h1q21)/+ cortex and hippocampus, and 72 and 48 in in Df(h15q13)/+ cortex and hippocampus), whereas the 15q13.3 homozygous samples displayed a much larger number of DE genes in both the cortex and hippocampus (476 and 564, respectively) (Figure 1A, Supplementary Figure 1A and supplementary table 2), consistent with its more pronounced behavioral and physiological phenotypes ^15^. Each CNV manifested a unique pattern of transcriptomic changes, and there was little overlap in differentially expressed (DE) genes across the CNVs (Figure 1B). Consistent with this, the top GO terms did not overlap between the different CNV/tissue combinations (Supplementary Figure 1B).

**Figure 1.**
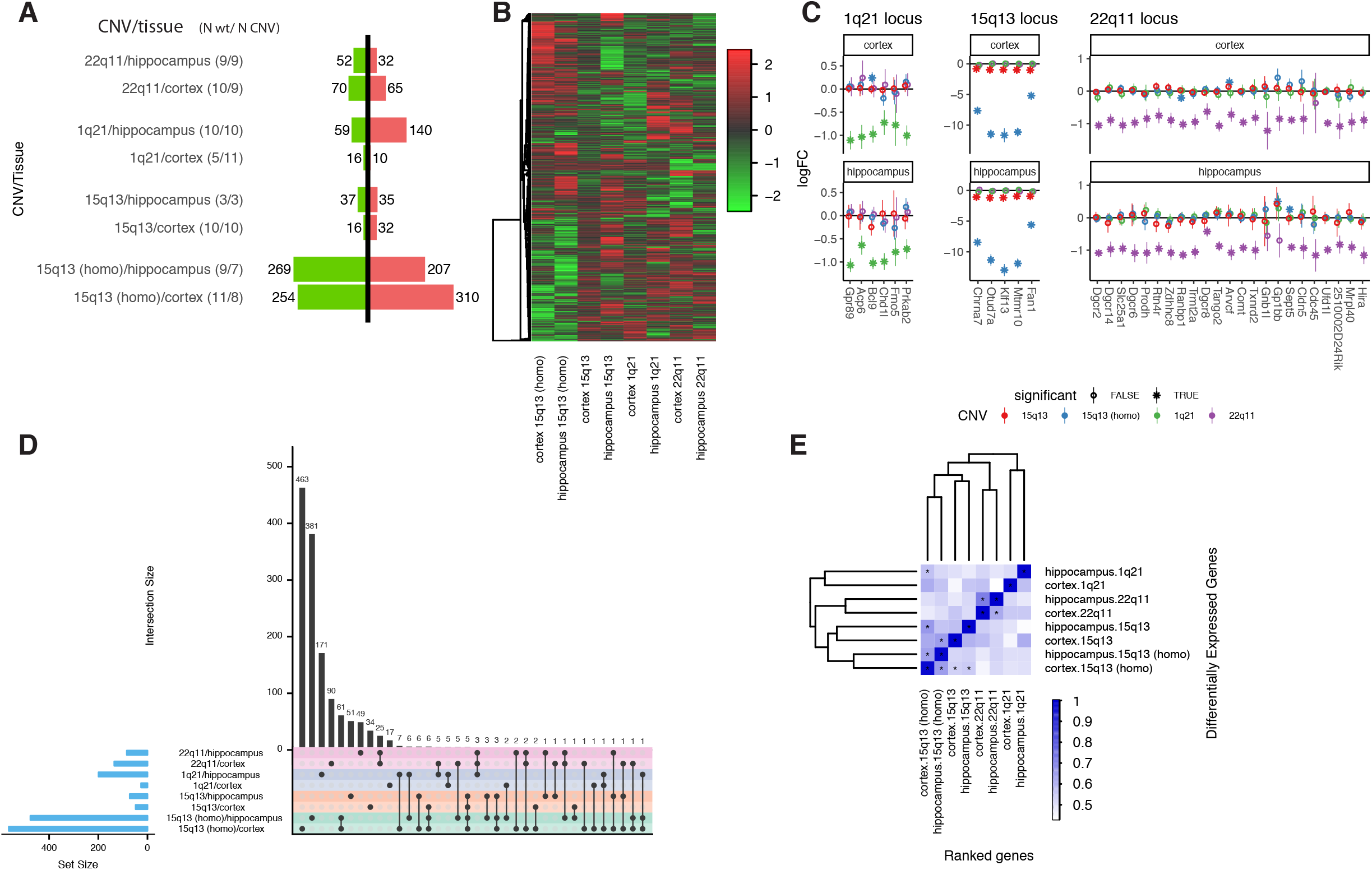
Differential gene expression in CNV mouse models of Psychiatric disorders. (A) Number of differentially expressed genes (DEG) in each CNV/tissue combination at p < 0.005. Parentheses are numbers of animals form each genotype after outlier removal (wt/mutant). Red and green bars denote up and down regulated genes, respectively. (B) A heat map of all differentially expressed genes in any of the CNV/tissue combinations. Values represent the scaled logFC. (C) Log fold change of the genes within the CNV loci. Color denote different CNVs while the shape represent significance at p < 0.005. (D) Overlap between Differentially expressed genes in the different CNV/Tissue combinations. An UpSet plot depicting each set in a row and the column correspond to intersections of the different sets. (E) Heatmap of mean gene rank. Mean gene rank scales from 0 (all non-differentially expressed genes are at the bottom of the gene list ranked by p value) to 1 (all differentially expressed genes are at the top of the gene list ranked by p value) p-values: * FDR < 0.05.

Close examination of the CNVs showed that 90-100% of the expressed genes in each of these loci were significantly downregulated in the corresponding mice and most showed a dosage dependent reduction in expression in the hemizygous mice (logFC of approximately −1 which is equivalent to 50% reduction) (Figure 1C) as has been observed in humans with 22q11.2 deletion ^52^. In the homozygous, Df(h15q13)-/- mouse, gene expression was reduced by 97%- 99.9% essentially eliminating expression of these genes.

### Overlap between the differentially expressed genes across regions and between distinct CNV carriers

We next tested for overlap of genes identified as DE at the single gene level between the hippocampus and cortex across the various CNV combinations and between these two brain regions within CNV. The highest overlap was observed between the different brain regions carrying the same CNV (Figure 1D). But, even in this case, only a minority of the genes overlapped. Defining DE genes requires the use of a (arbitrary) p-value cutoff. To reduce the effect this arbitrary cut off when comparing the DE genes, we also compared the mean ranking of DE genes (Methods). This allowed us to compare a list of DE genes from one condition to the entire set of genes from a different condition. We observed significant overlap between the hippocampus and cortex of the hemi- and homozygous Df(h15q13) mice, and between the different regions in the Df(h22q11)/- mice (Figure 1E). This was also evident by clustering the samples based on the mean effect size rank, where we observed that different tissues from the same CNV clustered together (Figure 1E). Using this analysis, we observed almost no overlap between mice harboring the different CNVs (Figure 1E).

### Comparisons with human post mortem gene expression changes in humans

We next wanted to compare the changes in gene expression identified in the different mouse models to those seen in the individuals with different psychiatric disorders. To this end, we compared the DE genes in the cortex of these mice to genes DE in the post mortem cortex of patients with different psychiatric disorders: autism spectrum disorder (ASD), schizophrenia (SCZ), bipolar disorder (BD), and major depressive disorder (MDD) alcohol abuse and dependence (AAD), including irritable bowel disease (IBD) as a control. Remarkably, we observe a significant overlap between downregulated genes in the Df(h15q13)−/− mouse cortex and downregulated genes in the cortex from individuals with ASD (OR = 1.7, fdr = 2e-05) and SCZ (OR = 1.4, fdr = 0.006). But, we could not detect any significantly overlap in the downregulated genes from any of the hemizygous mice (Figure 2A).

**Figure 2.**
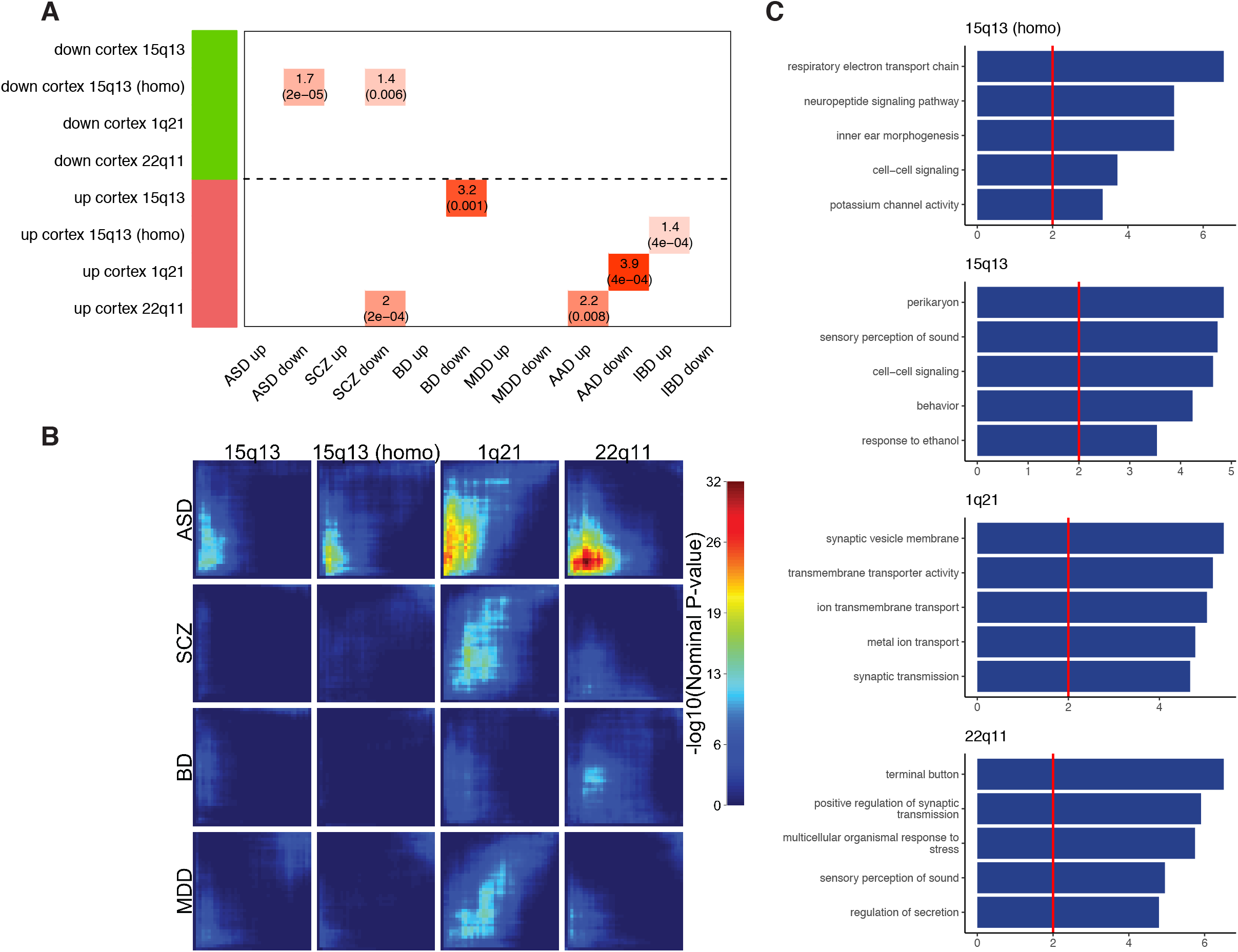
Comparison to human postmortem gene expression. (A) Enrichment of up and down regulated differentially expressed genes from different neurodevelopmental and psychiatric disorders from Gandal et al. (2018). (B) Rank-rank hyper geometric overlaps between the logFC of the cortical data from different mice and the logFC from different neurodevelopmental and psychiatric disorders from Gandal et al. (2018). (C) The top hub genes of the down regulated genes overlapping between ASD and the different CNVs in the RRHO (bottom left hand corner of the top row of B).

Since using an arbitrary p-value cut off to compare lists of genes results in loss of information and is dependent on power, we also performed a rank-rank hypergeometric test ^38^ which utilizes a sliding scale on both test and reference gene lists. Using this more sensitive test that comes closer to assessing genes as a group, we observed a significant overlap between genes down regulated in the cortex of the different mouse models in our study and genes down-regulated in the cortex of patients with ASD (Df(h15q13)−/+ p=6.98e-15, Df(h15q13)−/− p=7.45e-21, Df(h1q21)/+ p=1.73e-25, and Df(h22q11)/+ p=9.71e-33; bottom left corner of each plot in Figure 2B). These down regulated genes were all associated with GO terms related to neuronal function (Figure 2C). This finding is consistent with previous finding showing that the genes down regulated in ASD and SCZ are generally associated with neuronal function^4, 6^.

### Network analysis identifies gene modules shared across CNVs

We reasoned that a gene network approach, by assessing co-regulated, functional groups of genes, rather than analysis of individual genes as is the case with DE, would be a more powerful method to detect convergence between the different mouse models ^53^. To this end, we constructed a region specific co-expression network using Weighted Gene Co-expression Network Analysis (WGCNA)^36^. We removed CNV region specific effects by regressing out the CNV along with technical sequencing covariates (Methods), while retaining the effect of genotype (wt/hemizygous). As transcription was much more affected in the homozygous Df(h15q13)-/- causing them to be relative outliers (Figure 1A), we only used the hemizygous and wt mice transcriptome to construct the network.

Analysis of the cortical samples (Figure 3A and Supplementary Table 3) revealed two modules significantly associated with genotype (fdr < 0.1), which were regulated in the same direction across all CNVs: the up regulated cortical module 1 (cM1; adjusted R^2^ = 0.42, fdr < 0.1) and the down regulated cortical module 2 (cM2; adjusted R^2^ = 0.36, fdr < 0.1; Figure 3B). Examining the module eigengene of each sample for both of these modules showed that they were not driven by a single CNV, but rather showed similar dysregulated patterns across the all mouse models (Figure 3C).

**Figure 3.**
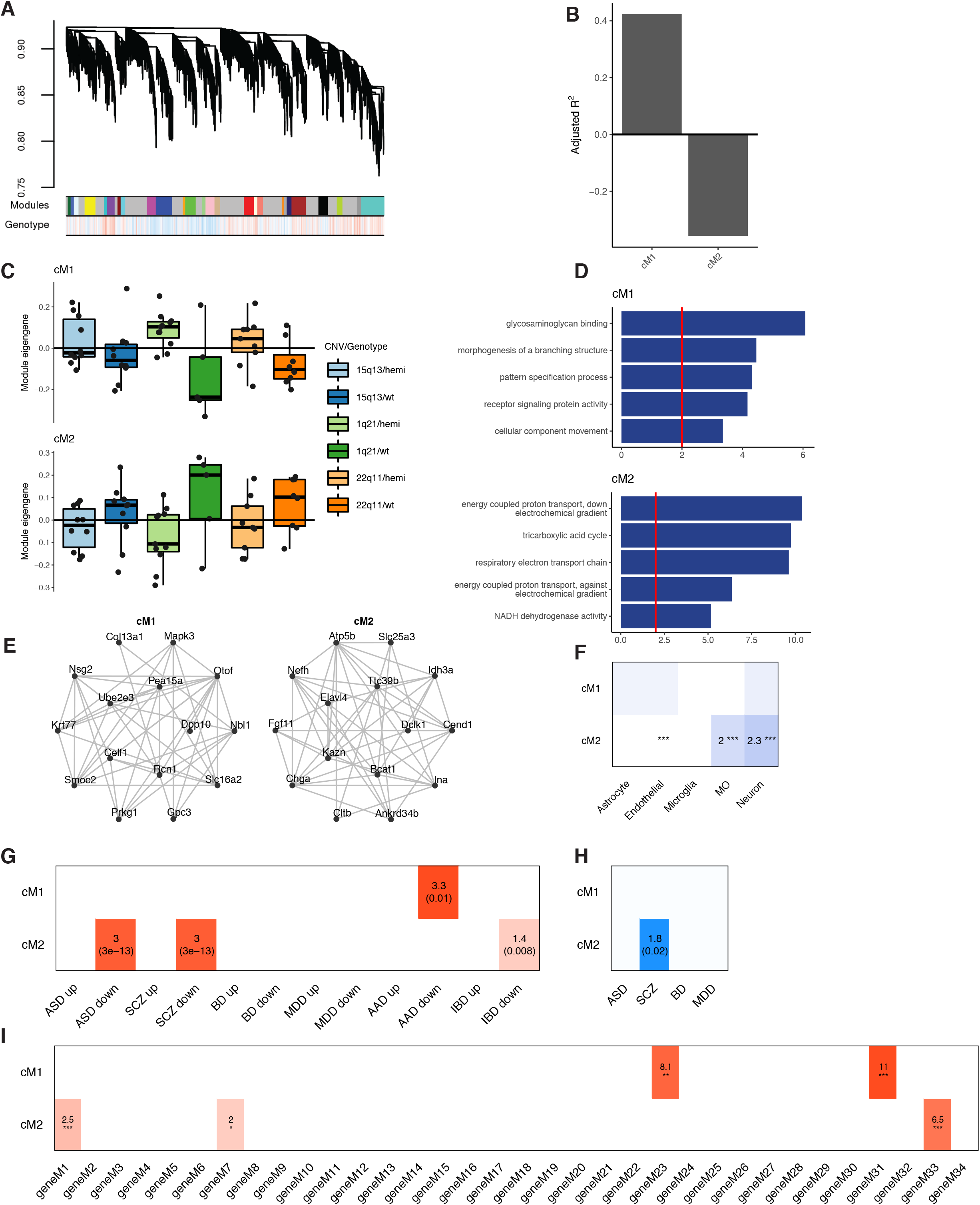
Network analysis of combined cortical data from the different CNVs. (A) Cortical network dendogram of genes across the different CNVs with module assignment. Color bar shows correlation to genotype. (B) Adjusted R-squared modules significantly associated with genotype across all CNVs with FDR < 0.1. (C) Module eigengene (ME) of modules significantly associated with genotype. (D) GO terms enriched in modules significantly associated with genotype. (E) Network plots of the top 15 hub genes with the top 50 connections between them for modules significantly associated with genotype. (F) CNS cell type enrichment. Values are fold enrichment. *** - fdr < 0.005 (G) Enrichment of Differentially expressed genes from different neurodevelopmental and psychiatric disorders from Gandal et al. (2018) in the modules significantly associated with genotype. Values are fold enrichments and fdr values are in parentheses. (H) Enrichment for GWAS for ASD ^44^, SCZ ^45^, BD ^1^, and MDD ^46^ using stratified LD score regression (FDR < 0.05). (I) Overlap of modules significantly associated with genotype with cross disorder genes modules ^5^. fdr * < 0.05, ** < 0.01, *** < 0.005. MO: myelinating oligodendrocyte.

To characterize these modules, we performed GO term enrichment (Figure 3E). The upregulated cM1 was enriched for genes that were linked to cell morphology (i.e. “morphogenesis of branching structures” and “pattern specification process”), while the downregulated cM2 was enriched for genes linked to mitochondrial related energy balance (i.e. “energy coupled proton transport” and “respiratory electron transport chain”; e.g. hub genes Slc25a3, Idh3a, Atp5b; Figure 3D). Testing for cell type enrichment revealed that while cM1 was not enriched for genes from a single cell type, cM2 was significantly enriched for neuronal genes (OR = 2.3, fdr < 0.005) and myelinating oligodendrocyte genes (OR = 2, fdr < 0.005; Figure 3F).

To test whether these modules were relevant to human disease, we measured whether they were enriched for dysregulated genes in post mortem brains from patients with psychiatric disorders ^4^. The upregulated cM1 was only enriched for genes upregulated in AAD^4^ (OR = 3.3, fdr = 0.01). Conversely, the downregulated cM2 was significantly enriched for genes down-regulated in ASD^4^ (OR = 3, fdr = 3e-13), SCZ^4^ (OR = 3, fdr = 3e-13), and to a lesser extent, from colon biopsy taken from patients with IBD^4^ (OR = 1.4, fdr = 0.008) (Figure 3G).

Since these modules were constructed from data coming from models of rare variation associated with SCZ, we reasoned that it would be important to know whether they were enriched for common variation associated with psychiatric disorders. We tested for enrichment of SNP heritability on the basis of GWAS results from ASD ^44^, SCZ ^45^, BD ^1^, and MDD ^46^ and found significant enrichment in cM2 for SCZ GWAS results (enrichment = 1.8, fdr = 0.02). We next compared our modules to those found in a psychiatric cross disorder transcriptomic study ^5^. cM1 was enriched for geneM23 (OR = 8.1, fdr < 0.01) and geneM31 (OR = 11, fdr < 0.005; Figure 3I), which is a module upregulated across ASD, SCZ and BD ^5^. cM2 overlapped with gene modules geneM1 (OR = 2.5, fdr < 0.005), geneM7 (OR = 2, fdr < 0.05) and geneM33 (OR =6.5, fdr < 0.005) (Figure 3I), both neuronal modules enriched for mitochondrial processes. Interestingly, the previously identified geneM7 was also enriched for both SCZ and BD GWAS signals ^5^.

The hippocampal co-expression network (Supplementary Figure 2A and supplementary table 3) identified 10 modules that were significantly associated with genotype (Supplementary Figure 2B of which two modules were dysregulated in the same direction in all CNVs (Supplementary Figure 2C). hM1 was most enriched for unfolded protein binding, whereas hM2 module was most enriched for nucleosome organization (Supplementary Figure 2E). Comparing the hippocampal and cortical modules revealed that both hippocampal modules significantly associated with genotype also showed significant overlap with cortical modules; cM1 overlapped with the hM7 module (OR = 4.4, fdr = 1.67e-02) and cM2 overlapped with both hM4 and hM5 (OR = 3.2, fdr = 6.69e-5 and OR = 2.3, fdr = 2.85e-2 respectively; Supplementary Figure 3A).

To test whether the modules identified replicated in another data set, we conducted preservation analysis in the Df(15q13)-/- homozygous mice, which were not used to construct the network. In the cortex, cM2 was very strongly preserved (Z_summary_= 19) while cM1 was moderately preserved (Z_summary_= 7.2; Supplementary Figure 3B). Both of these modules were dysregulated in the same direction in the Df(15q13)-/- samples (Supplementary Figure 3C). In the hippocampus, 7 of the 10 modules were strongly preserved, including both hM1 (Z_summary_= 12) and hM2 (Z_summary_= 21; Supplementary Figure 3B). All but two of these modules (hM6 and hM7) were dysregulated in the same direction in the Df(15q13)-/- samples (Supplementary Figure 3C), consistent with substantial overall preservation.

### Characterization of the downregulated cortical mitochondrial module

We further characterized the down-regulated cM2, which was the only module that was preserved across both regions in all transgenic strains and overlapped with patterns observed in SCZ and ASD post mortem brain transcriptomic changes. We first elaborated on the mitochondrial signature found in this module. cM2 was significantly enriched for both classes of transcriptionally distinct mitochondria found in neurons^54^: synaptic mitochondria (OR = 4.79, fdr = 9e-10) and soma enriched neuronal activity-related mitochondria (OR = 5.03, fdr = 1.13e-14; Figure 4A). cM2 also significantly overlapped with a neuronal mitochondrial module found to be down-regulated in comprehensive RNA sequencing performed in post-mortem tissue from multiple psychiatric disorders (ASD, SCZ, BD) ^4^ (OR = 2.72, fdr = 1.33e-8; Figure 4A), further validating its disease relevance and potential generalizability of genes within this module.

**Figure 4.**
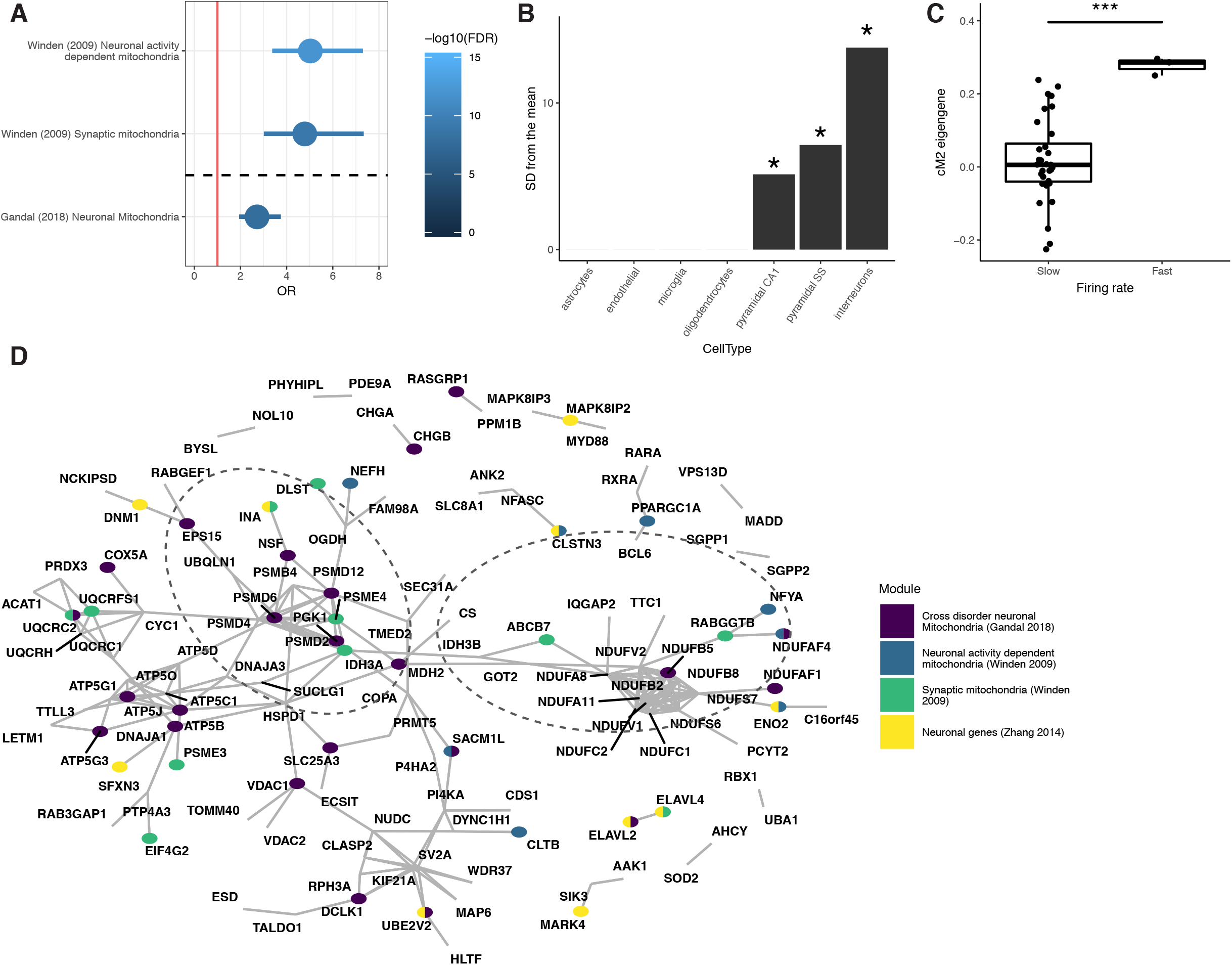
cM2 overlap with mitochondria subtypes. (A) Significant overlap of cM2 with two neuronal mitochondrial types from Winden et al. 2009 OR = 5.03 (35/396 genes overlapping) for soma enriched neuronal activity-related mitochondria and OR = 4.7 (22/396 genes overlapping) for synaptic mitochondria as well as with the mitochondrial module from Gandal et al. 2018 (OR = 2.7 51/269 genes). (B) Cell type enrichment of cM2 using mouse cortical single cell ^42^. (C) Box plot of cM2 eigengene in fast and slow firing neurons ^47^. *** p < 0.005 Mann-Whitney-Wilcoxon test. (D) Protein-Protein interaction network. The colors represent overlap with the above modules. * p < 0.05.

To obtain a more detailed picture of which cell types were represented by this module, we next performed Expression Weighted Cell-type Enrichment (EWCE)^41^ using single cell expression data from mouse cortex and hippocampus ^42^. We observed the largest enrichment in inhibitory neurons (SD away from mean = 13.7 and 7.1, fdr < 0.005) with less, but still significant, enrichment in excitatory neurons (SD away from mean = 5; Figure 4B and Supplementary Figure 4). Due to the association of neuronal firing rate with cellular energetics and mitochondrial genes, we hypothesized that this module was related to neuronal firing rate^54^. We calculated the module eigengene for the different neuron types ^47^ and also classified the neurons based on the firing rate in current clamp measurements from the same study ^47^. We observed a significant increase in the expression of cM2 genes (as measured by the module eigengene) in fast firing neurons compared to slow firing neurons (p < 0.005; Figure 4C) ^54^.

To further validate this module as representing a biologically coherent process, we analyzed the protein-protein interactions (PPI) of the genes found in this module. The resulting network showed a significant enrichment for PPI (p = 0.001) demonstrating that this module represents a biological relevant functional unit. The PPI network structure revealed two functional clusters within this module, a mitochondrial cluster containing many NDUF proteins and a proteasome-mitochondrial related cluster containing many PSM proteins (Figure 4D) further confirming the biological relevance of this module.

## Discussion

Although studies on individual mouse models of psychiatric disorders, or single brain regions have been conducted ^9, 55–58^, no study has performed a comprehensive comparison across several models to assess convergence. In this study we aimed to determine whether the patterns of changes in gene expression in the brains of mice carrying CNVs that increase risk for SCZ and other psychiatric disorders were distinct or overlapping. We found that each CNV results in a unique pattern of DE genes not only between CNVs, but also between different brain regions within the same animal model. Despite these differences, network analysis allowed us to identify a key point of convergence: two gene modules that were dysregulated in all CNVs in both the cortex and hippocampus. One of these dysregulated modules, the neuronal mitochondrial module cM2, overlaps with changes previously observed in SCZ and ASD in human post mortem brain^59–61^, highlighting its broader relevance to humans with these disorders and not only to those harboring these specific CNVs. The observation of this same biological process previously observed in human brain in three distinct mouse models is significant because it eliminates the explanation that the neuronal energetic dysfunction observed in humans reflects a post mortem artifact ^62, 63^. Here, the well-controlled experimental design, lack of antemortem illness, absence of drug treatments, rapid processing of tissue, and high RNA integrity control for confounders that confront interpretation of post mortem human studies. Rather, the GWAS enrichment and the absence of the aforementioned confounders in the current study suggest a role for this process in SCZ.

### Face validity

We recognize that no single model is a replica of a human psychiatric disorder. At the same time, the relation between the CNV and potential human disease-related phenotypes in these models, so called “face validity,” is of interest ^10^. As previously shown, there is little similarity between the disease relevant behaviors in the three mouse models studied ^9^. This is consistent with the individuals carrying these CNVs ^64^, as well as the gene expression patterns observed in this study.

At the gene expression level, we found that the GO terms associated with DE genes identified in this study were often related to previous findings in patients with these CNVs. For example in the Df(h22q11)/+ mice the DE genes were enriched for calcium ion related GO terms (“calcium ion binding” and “cellular response to calcium ion”) a finding that has been replicated in another study^65^ and may be linked to the hypocalcemia seen in many 22q11 deletion carriers ^66^. Another example is the GO term associated with histone modifiers (“histone deacetylase binding”), which was enriched in the Df(h22q11)/+ mouse cortex and have been linked to the pathophysiology seen in patients with a 22q11.2 deletion ^67^.

### Convergence on mitochondria between CNVs

Gene network analyses, which considers patterns of co-regulated genes ^36^, revealed two cortical and hippocampal modules that were dysregulated in all three models. cM2 is of special interest, as it was preserved and changed in the same direction in ASD and SCZ post mortem brain, supporting its broader relevance to ASD and SCZ. cM2 showed a significant enrichment for inhibitory and, to a lesser extent, excitatory neuronal genes as well as for GO-terms related to mitochondrial activity (i.e. “energy coupled proton transport”). Moreover, the top hub genes of cM2 contain primarily neuronal (Nefh, Elavl4, Dclk, Cend1, Ina, and Chga) and mitochondrial (Atp5b, Slc25a3, Bcat1, Idh3a) genes. cM2 also significantly overlapped with neuronal-activity dependent mitochondrial and synaptic mitochondrial modules that were previously defined in mouse^54^, as well as with a neuronal mitochondria module that is down-regulated in post mortem brain from patients with ASD and SCZ ^4, 5^. These enrichments together with the increase in the expression of cM2 in fast firing neurons lead us to annotate this module as a neuronal-associated mitochondrial, or neuronal energetics module associated with psychiatric disorders.

In neurons, mitochondria play multiple roles including producing the energy that is required to maintain the resting membrane potential, axonal transport and vesicle recycling and docking ^68^, as well as regulating calcium concentration in dendritic spines^68^. In SCZ, it has been suggested that mitochondrial dysfunction impairs energy balance, as well as the ability of neurons and glia to resist and adapt to environmental stressors and their ability to undergo synaptic changes associated with plasticity ^69^. The increase of cM2 gene expression in fast firing neurons suggests that this module is linked to energy production required for maintaining rapid neuronal firing rates that are most prominent in certain classes of interneurons.

The identification of a neuronal energetics module corroborates multiple lines of genetic ^59^, metabolic ^60^, anatomic ^61^ and functional ^70^ evidence linking mitochondrial dysfunction to SCZ. Post-mortem brain studies show that mitochondrial structure and function are altered in SCZ, including a reduction in the number of mitochondria in neurons ^27, 61^. The association of SCZ and mitochondrial function is also supported by 28 mitochondrial related genes being contained among the 108 GWAS loci for SCZ, ^71–73^ which is consistent with our finding of enrichment of SCZ GWAS signals in cM2. Combined, these multiple lines of evidence, suggest a causal role for this pathway in the pathophysiology of SCZ.

The overlap of the genes in this neuronal mitochondrial module with down-regulated genes in both SCZ and ASD suggests that mitochondrial dysfunction is not unique to SCZ, but rather may be shared across multiple psychiatric disorders. This is in line with the pleiotropic nature of these CNVS ^19, 74^ and is concordant with emerging evidence linking mitochondria dysfunction in neurons across several psychiatric disorders ^4, 69^. Consistent with this, recent mega analyses of postmortem brains across six psychiatric disorders found a neuronal mitochondrial module that was downregulated in ASD, SCZ and bipolar disorder ^4, 5^, which highly overlaps with the mitochondrial module identified in this study.

Our results link mitochondria across the studied CNV models, and more broadly across relevant human neuropsychiatric disorders even without these CNV, but they do not identify the mechanism by which dysfunctional neuronal energetics are associated with SCZ or ASD. Our results suggest that these mouse models could be used to further assess the role and relationship of mitochondrial dysregulation and neuronal activity in cognitive and behavioral phenotypes. In addition, it would be especially interesting to test whether additional CNVs associated with SCZ and ASD ^2^ also converge onto neuronal firing or mitochondrial dysfunction, and whether normalizing this function could ameliorate disease-related phenotypes.

## Supplementary data

**Supplementary Table 1**– Sample meta data. (xlsx file)

**Supplementary Table 2**– Details of differential expression of all genes in the different CNVs/tissues. (xlsx file)

**Supplementary Table 3**– Module assignment and kMEs for all genes in both cortex and hippocampus. (xlsx file)

**Figure S1.**
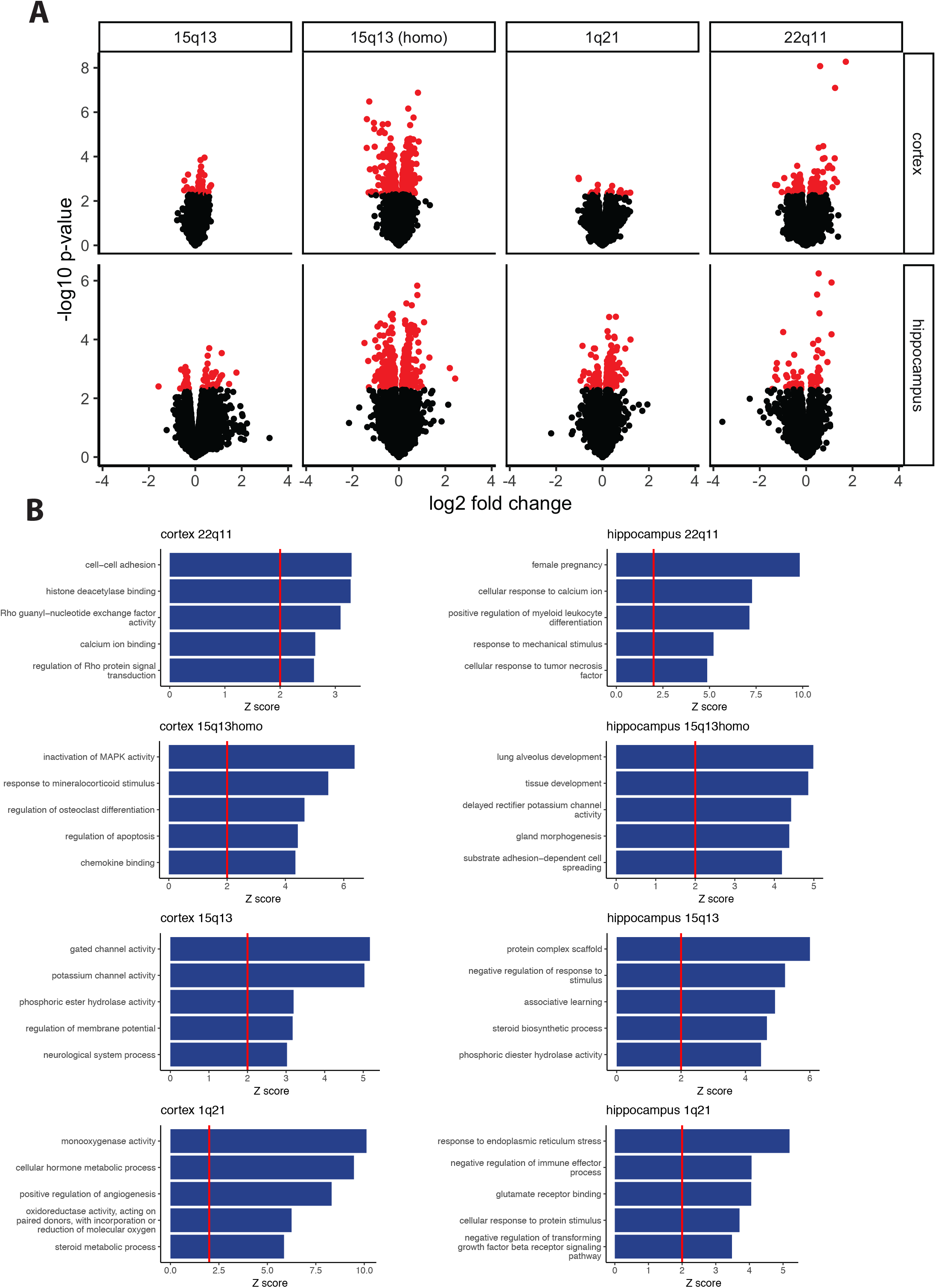
Effect size and GO term enrichment of DEG. (A) Volcano plots of DEG in tehe different CNVs and tissues. Red points represent genes with p value < 0.005. (B)Top 5 significant GO terms enriched in each CNV/tissue combination.

**Figure S2.**
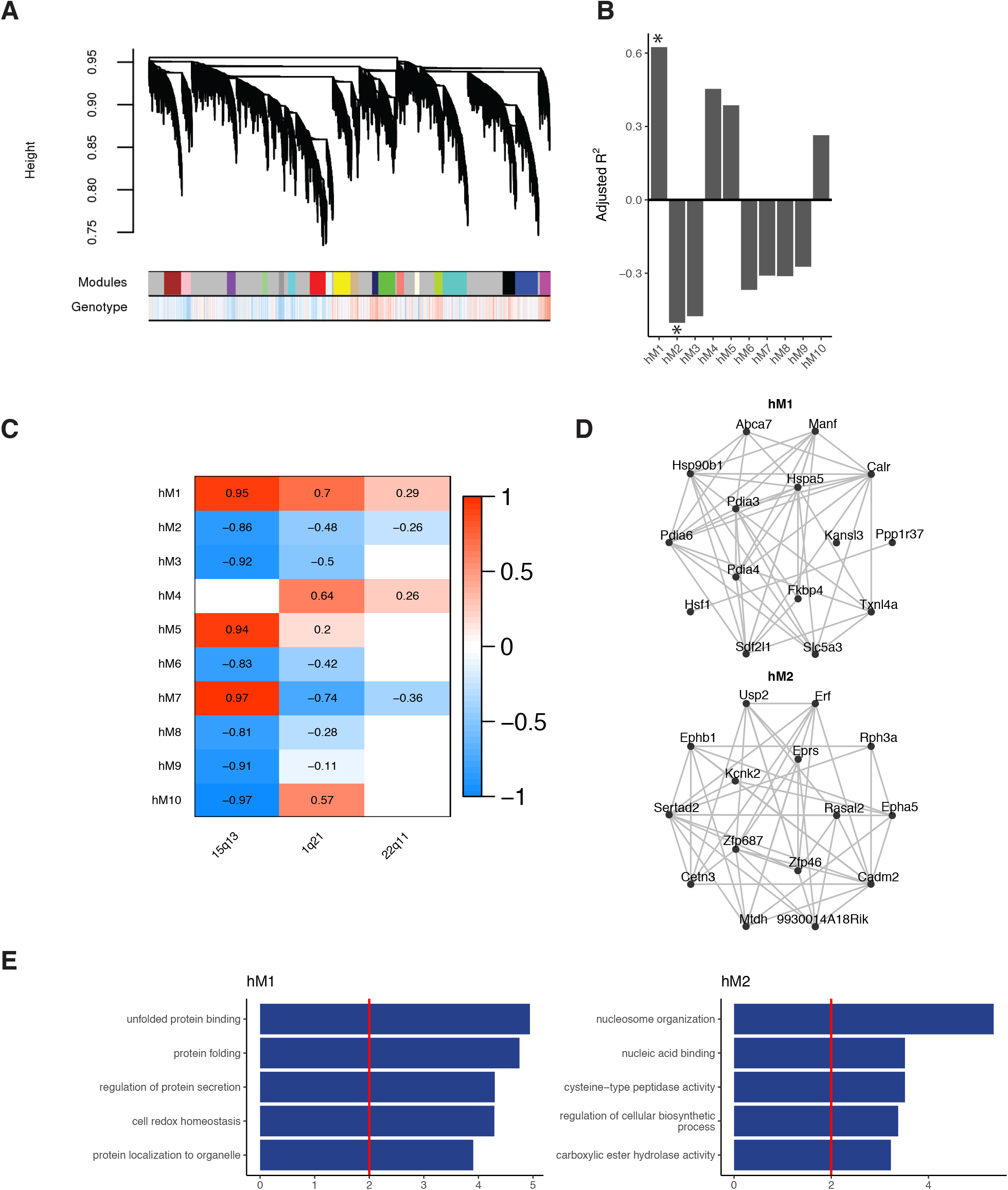
Network analysis of combined hippocampal data from the different CNVs. (A) Hippocampal network dendrogram of genes across the different CNVs with module assignment. Color bar shows correlation to genotype. (B) Adjusted R-squared of modules significantly associated with genotype. Asterisk denote modules with the same direction association in all CNVs. (C) Adjusted r-squared of modules with each CNV separately. (D) Network plots of the top 15 hub genes with the top 50 connections between them for modules significantly associated with genotype. (E) GO terms enriched in modules significantly associated with genotype.

**Figure S3.**
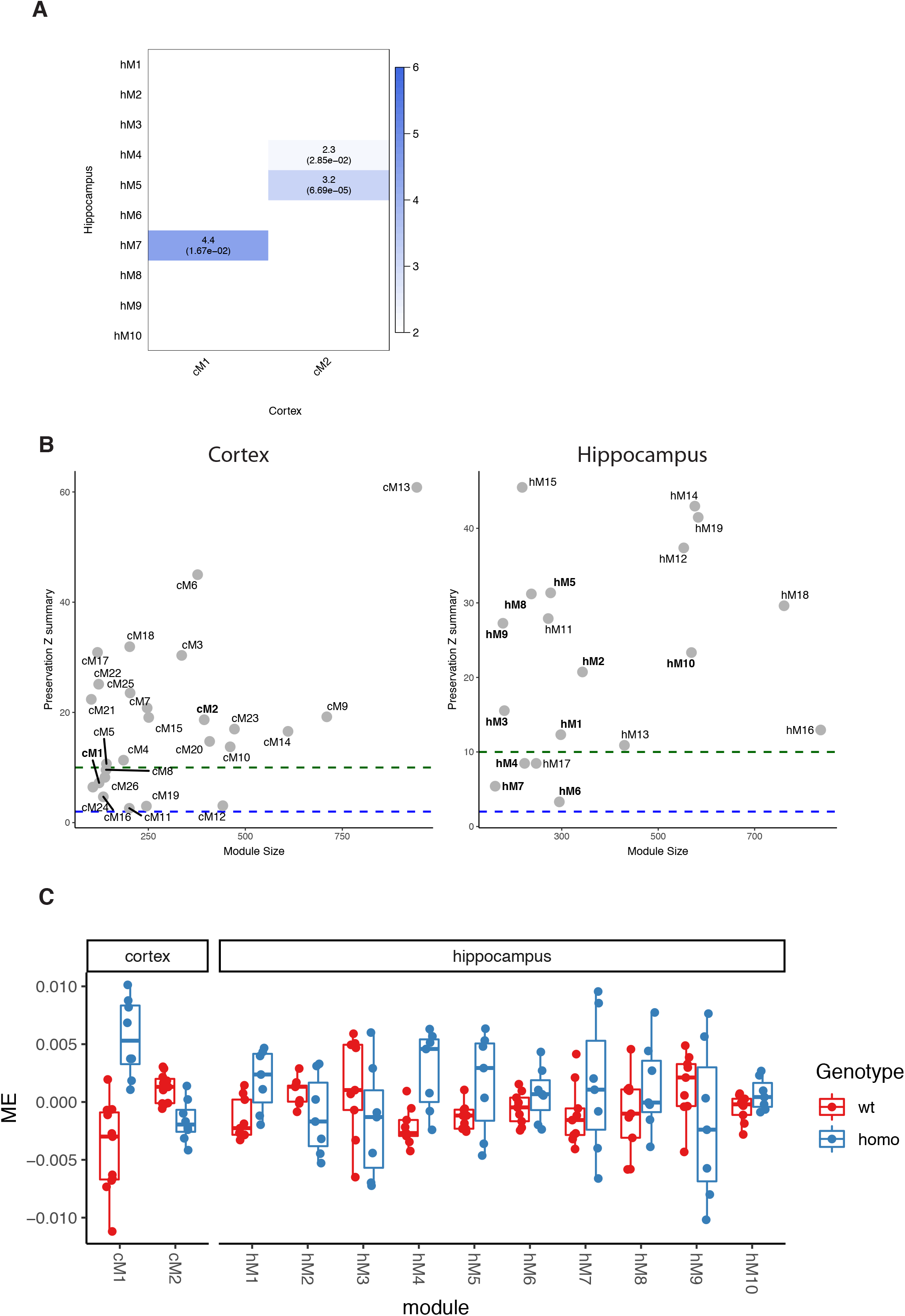
Cortical and hippocampal module overlap and preservation. (A) Overlap of genes in each module. OR and fdr (in parentheses) are indicated. Color denotes OR. (B) Module preservation of the cortical (left) and hippocampal (right) modules in the 15q13 homozygous samples. Modules above Z_summary_ > 10 are considered to have strong evidence for preservation while modules with Z_summary_ > 3 are considered to have moderate evidence for preservation. (C) Module eigengenes (first principal component the module) of the Df(h15q13)−/− samples.

**Figure S4.**
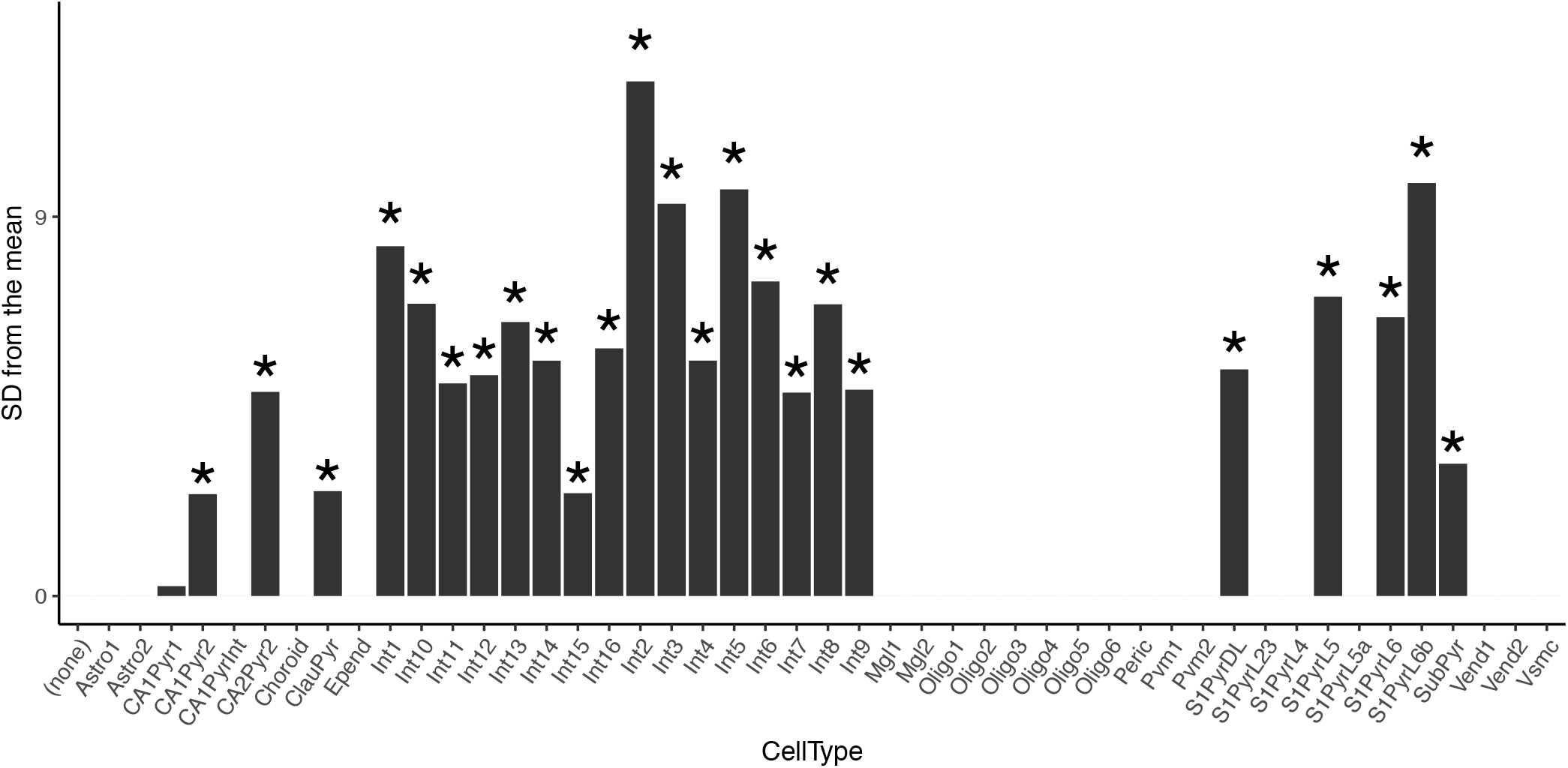
Detailed cell type enrichment using Expression Weighted Cell-type Enrichment. Enrichment of cM2 genes in different mouse cortical and hippocampal cell types ^42^ using EWCE ^41^. * - fdr < 0.05.

